# An optogenetic method for the controlled release of single molecules

**DOI:** 10.1101/2023.09.16.557871

**Authors:** Purba Kashyap, Sara Bertelli, Fakun Cao, Yulia Kostritskaia, Fenja Blank, Niranjan Srikanth, Roberto Saleppico, Dolf Bierhuizen, Xiaocen Lu, Walter Nickel, Robert E. Campbell, Andrew Plested, Tobias Stauber, Marcus J. Taylor, Helge Ewers

## Abstract

We developed a system for optogenetic release of single molecules in live cells. We confined soluble and transmembrane proteins to the Golgi apparatus via a photocleavable protein and released them by short pulses of light. Our method allows for the controlled delivery of functional proteins to cytosol and plasma membrane in amounts compatible with single molecule imaging, greatly simplifying access to single molecule microscopy of any protein in live cells. Furthermore, we could reconstitute cellular functions such as ion conductance by delivering BK and VRAC ion channels to the plasma membrane. Finally, we could induce NF-kB signaling in T-Lymphoblasts stimulated by IL-1 by controlled release of a signaling protein that had been knocked-out in the same cells. We observed light induced formation of functional inflammatory signaling complexes that could trigger IKK phosphorylation in single cells. We thus developed an optogenetic method for the reconstitution and investigation of cellular function at the single molecule level.

---

Single molecule fluorescence microscopy is a powerful technique for the investigation of protein function. Many fundamental cellular processes, including stepping of motor molecules^1^, DNA replication^2^, transcription^3,4^ and translation^5^ or the stoichiometry^6^ and mechanism of action^7^ of membrane receptors have been elucidated using single molecule imaging. However, single molecule imaging requires a sparse population of labeled molecules to avoid overlap of signal. At the same time, it is essential that no unlabeled endogenous background population interferes with the measurement, which is challenging to achieve. While genome engineering has alleviated the problem of quantitative and endogenous labeling, the controlled delivery of single molecules remains an unsolved challenge as existing approaches are either too leaky for control at the single molecule level since they are based on noncovalent attachment or require complex expression systems such as mRNA injection.

Here we have developed a technique for the optogenetically-controlled release of functional single molecules in live cells. We make use of a recently developed green to red photoconvertible protein called PhoCl^8^ that is photocleaved, splitting into two pieces after activation in near UV-light (Fig. 1a). In PhoCl, the fluorophore encoded by the amino acid sequence breaks upon UV-illumination, severing the peptide chain. Since PhoCl is circularly permutated, the fluorophore is located towards the end of the coding sequence. On breakage, the very C-terminus thus dissociates from the protein barrel structure. Here, we sequestered target proteins to the Golgi apparatus by fusing their coding sequence via PhoCl to Golgi resident proteins. This system allowed us to release functional target molecules at amounts compatible with single molecule imaging in a light controlled manner. At the same time the fusion to a Golgi-resident protein allowed for covalent attachment to the Golgi apparatus, resulting in low leakiness. We demonstrate the function of our system with the examples of a cytosolic protein, a single-spanning transmembrane protein, multi-subunit ion channels and a kinase within a signaling complex. Our system allowed controlled single molecule imaging at the plasma membrane and the surface delivery of single transmembrane proteins. Furthermore, we could reconstitute specific ion channel conductance and restore a signaling pathway in knockout cells by releasing an essential kinase and subunit of the Myddosome signaling complex. We provide a new experimental paradigm that will allow both for quantitative single molecule imaging as well as optogenetic control over membrane protein function.

**Figure 1.**
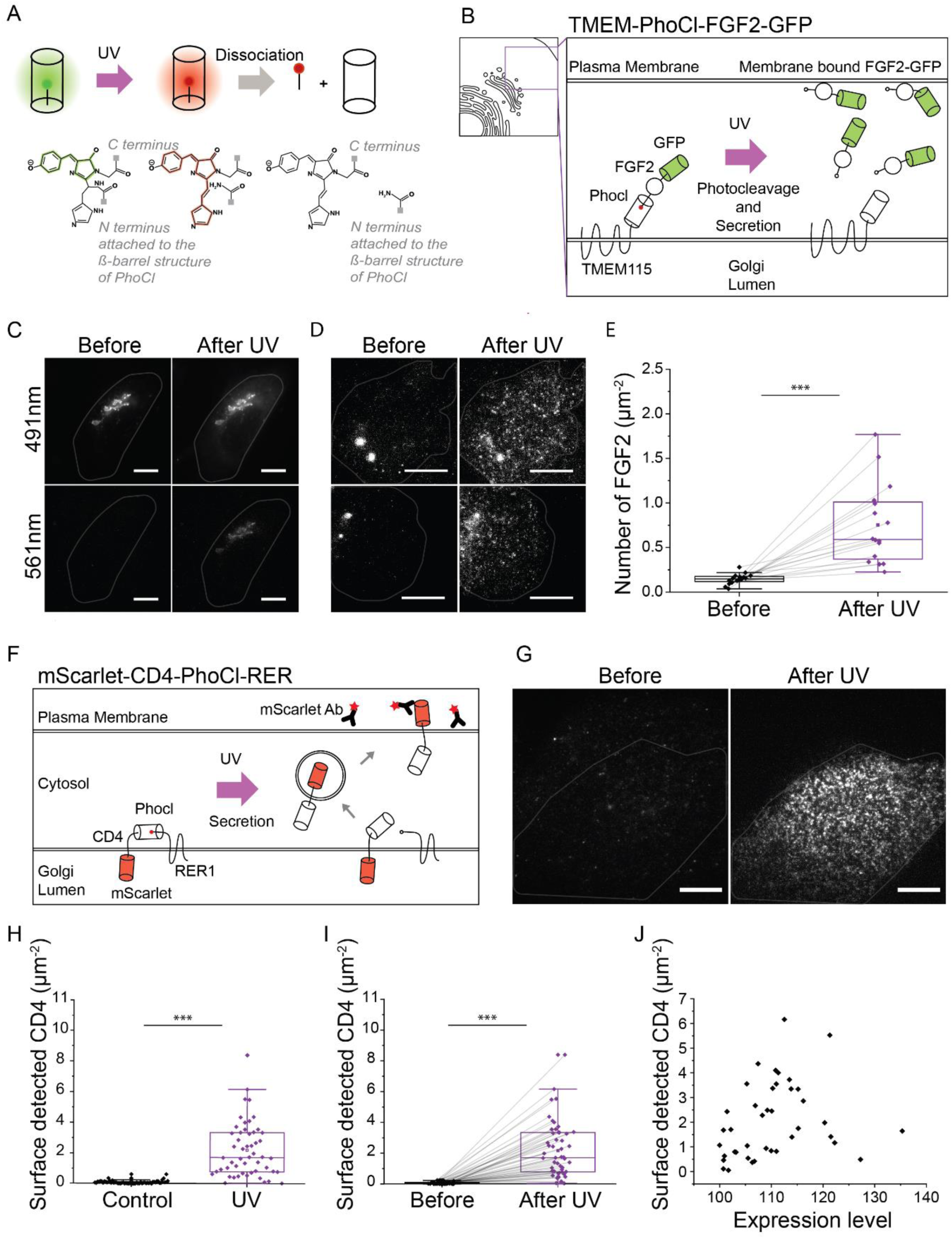
PhoCl based uncaging of cytosolic and transmembrane protein. **(a)** Schematic of light induced photoconversion and dissociation of PhoCl. **(b)** Schematic of TMEM-PhoCl-FGF2-GFP construct. **(c)** Images of a cell expressing TMEM-PhoCl-FGF2-GFP in 491 nm and 561 nm channels before and after UV illumination. **(d)** TIRF images of cells expressing TMEM-PhoCl-FGF2-GFP before and after UV illumination. **(e)** Quantification of membrane bound FGF2-GFP before (mean, sd = 0.14 ± 0.05) and after UV illumination (mean, sd = 0.75 ± 0.4), N = 2, n = 16. Significance was tested using the Paired sample sign test (p = 10^-4^). **(f)** Schematic of mScarlet-CD4-PhoCl-RER construct. Plasma membrane localized mScarlet-CD4 is recognized by labeled antibodies in the media. **(g)** TIRF images of a cell with antibodies against mScarlet before and after UV illumination. **(h)** Quantification of the number of anti-mScarlet antibodies bound to the control (mean, sd = 0.09 ± 0.1, n = 55) and UV illuminated cells (mean, sd = 2.20 ± 1.78, n = 52), N = 11. Significance was tested using the Mann-Whitney test (p = 10^-16^). **(i)** Quantification of the number of anti-mScarlet antibodies bound to the same cells before (mean, sd = 0.06 ± 0.07) and after uncaging (mean, sd = 2.16 ± 1.77), N = 11, n = 51. Significance was tested using the Paired sample sign test (p = 10^-9^). **(j)** Correlation of expression level of mScarlet-CD4-PhoCl-RER with the amount of secreted CD4 detected after uncaging, N = 11, n = 47. Spearman‘s correlation test, r_s_ = 0.26, p = 0.07. Scale bars are 10 µm. N = number of biological replicates, n = number of cells, sd = standard deviation. The center line in all the box plot represents the median, the box represent the 25%-75% of the data, whiskers represent 1.5 interquartile range.

## Results

We first asked if a soluble molecule could be confined to a specific location in the cell and released upon photocleavage of PhoCl. To do so, we made use of FGF2, a cytosolic protein that becomes secreted via an unconventional pathway through the plasma membrane^9,10^. Before penetrating the plasma membrane, FGF2 can be readily visualized in single molecule total internal reflection fluorescence (TIRF) microscopy at the inner membrane leaflet and upon secretion detected from outside of the cell^9^. To sequester FGF2 outside the evanescent field of TIRF, we generated a fusion construct from FGF2, PhoCl and the Golgi resident transmembrane protein TMEM115 (Fig. 1b). When we expressed this construct, we could observe FGF2-GFP at the Golgi apparatus (Fig. 1c). When we then illuminated the sample with brief pulses of 405 nm light, we could detect red fluorescence of PhoCl appear at the Golgi apparatus, besides persistent green fluorescence, indicating breakage of the peptide chain backbone in a fraction of molecules (Fig. 1c). Consistent with this, quantification before and 10 min after the activation showed an increase in overall cytosolic FGF2-GFP fluorescence (Supplementary Fig. 1). Soon after the UV-pulse, highly mobile single FGF2-GFP spots appeared in the TIRF field (Supplementary movie 1, Supplementary Fig. 2) suggesting recruitment to the inner plasma membrane leaflet (Fig. 1d, e). We concluded that FGF2-GFP sequestration to the Golgi apparatus as well as its optogenetic release were successful and yielded functional FGF2 that was recruited to the plasma membrane.

Next, we aimed to use our optogenetic method to release single transmembrane proteins. To do so, we used mScarlet-CD4, a generic, single-spanning plasma membrane protein. We created a construct containing the coding sequences of mScarlet-CD4, PhoCl and Retrieval Protein-1 (RER), a Golgi resident protein (Fig. 1f). RER was able to successfully anchor mScarlet-CD4 at the Golgi apparatus (Supplementary Fig. 3a). Like before, UV illumination of PhoCl led to uncaging and trafficking of mScarlet-CD4 to the plasma membrane, where multiple laterally mobile single molecule spots appeared in TIRF microscopy (Fig. 1g, Supplementary movie 2). We found that the amount of plasma membrane inserted molecules could be controlled by the strength of the uncaging pulse (data not shown), suggesting that the amount of released molecules can be controlled by light intensity. To ensure that only mScarlet-CD4 molecules properly inserted into the plasma membrane were quantified, we detected mScarlet-CD4 from the exoplasmic space. To do so, we added Alexa Fluor 647 (AF647) labelled antibodies to live cells and quantified them in TIRF microscopy before and after activation (Fig. 1h, i). When we quantified the number of surface detected molecules against the expression level before uncaging, we found that higher expression level led to more surface delivered molecules with the same pulse intensity (Fig. 1j). We concluded that PhoCl in combination with RER as a Golgi anchor can be used to optogenetically control the delivery of a transmembrane protein to the plasma membrane at levels compatible with single molecule imaging in a quantitative manner.

Next, we aimed to reconstitute functional ion channels into the plasma membrane of cells. To do so, we made use of the BK-channel, a voltage-gated and a calcium-sensing potassium channel^11^ that is assembled from four identical subunits. When we fused the coding sequence of the BK-channel to TMEM115 via PhoCl (Fig. 2a), we found that it localized to the Golgi apparatus after transient expression (Supplementary Fig. 3b). When we then illuminated cells with a 5s pulses of 405 nm light (33 mW/mm^2^), we found that surface-delivered BK-channels could be detected in TIRF microscopy via AF647-coupled antibodies added to the medium (Fig. 2b-d). In contrast, they were not detectable in the same cells without 405 nm illumination (Fig. 2b-d). Consistent with this, electrophysiological measurements after uncaging revealed a dramatic increase of characteristic BK-channel currents compared to control cells (Fig. 2e, f). However, a basal current could be recorded also for control cells which were not illuminated, indicating a minor leakage (Fig. 2e, f). We concluded that our system allows for the controlled release of functional ion channels and thus multispanning multisubunit transmembrane proteins.

**Figure 2.**
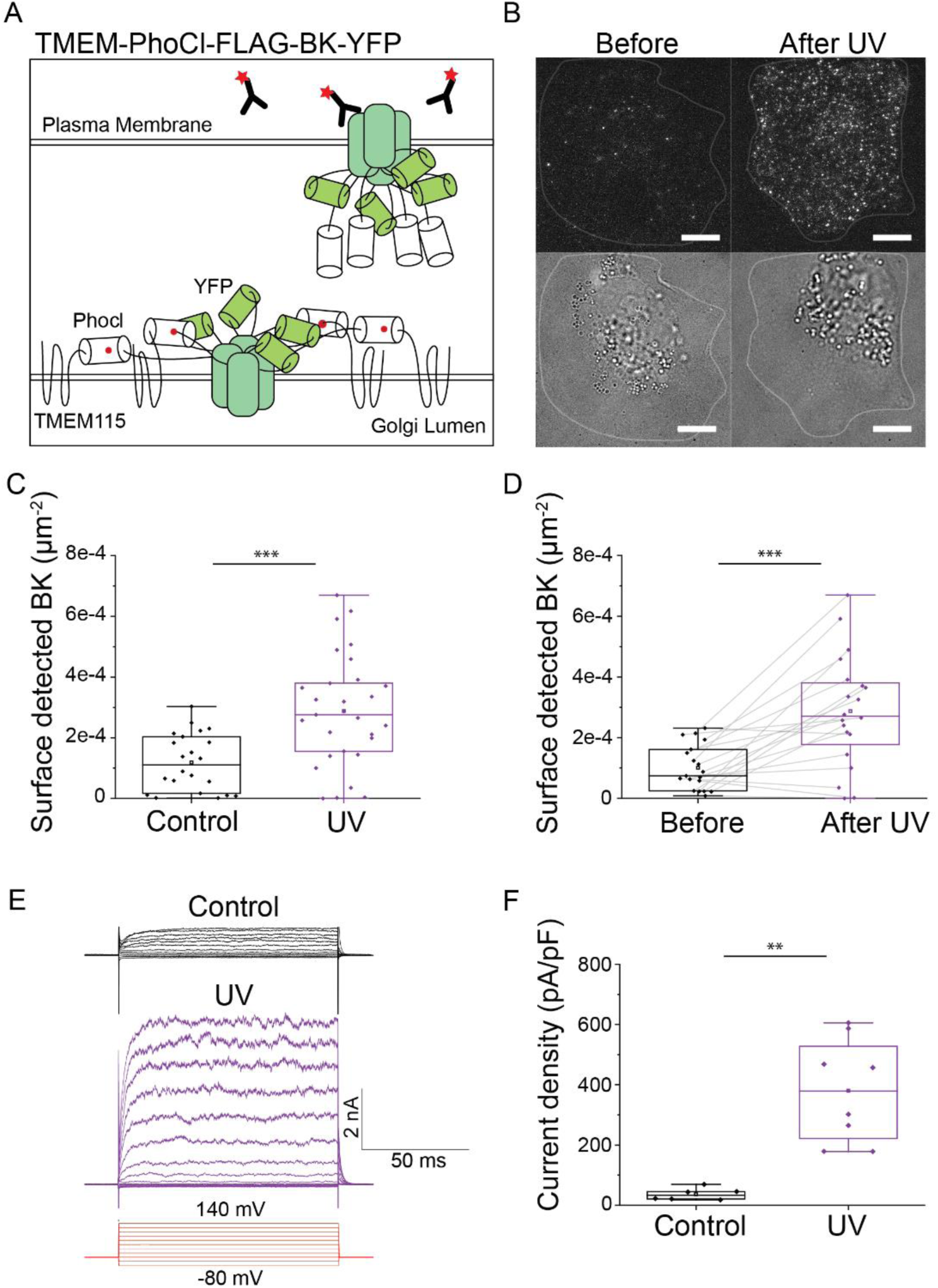
Optogenetic release of BK-channels. **(a)** Schematic of FLAG-BK-YFP-PhoCl-TMEM construct. Following uncaging, BK-channels secreted to the membrane were detected using antibodies in the media. **(b)** TIRF and brightfield images of a cell with antibodies against FLAG before and after UV illumination. **(c)** Quantification of the number of anti-FLAG antibodies bound to the control (mean, sd = 1.1e^-4^ ± 9e^-5^, n = 22) and UV illuminated cells (mean, sd = 2.8e^-4^ ± 1.8e^-4^, n = 29), N = 5. Significance was tested using the Mann-Whitney test (p = 10^-4^). **(d)** Quantification of the number of anti-FLAG antibodies bound to the same cells before (mean, sd = 1e^-4^ ± 7.3e^-5^) and after uncaging (mean, sd = 2.8e^-4^ ± 1.8e^-4^), N = 5, n = 19. Significance was tested using the Paired Wilcoxon signed rank test (p = 0.001). **(e)** Representative current traces of BK-channel at voltages with the protocol shown in red in control and UV illuminated cell. **(f)** Quantification of current density at 120 mV in control (mean, sd = 36 ± 20, n = 6) and UV illuminated cell (mean, sd = 380 ± 172, n = 8), N = 4. Significance was tested using the Mann-Whitney test (p = 0.002). Scale bars are 10 µm.

Encouraged by these results, we decided to reconstitute an ion channel with more complex regulation. We chose <u>V</u>olume-<u>R</u>egulated <u>A</u>nion <u>C</u>hannels (VRACs) that are formed by LRRC8 proteins. VRACs open upon osmotic cell swelling and exhibit electrophysiological properties defined by the subunit composition^12^. The hexameric VRACs are constituted from at least one LRRC8A subunit that is essential for export from the endoplasmic reticulum and surface delivery, and heteromerizes with at least one of the LRRC8B-E subunits^13^ (Fig. 3a). We expressed LRRC8E-GFP and LRRC8A-PhoCl-TMEM in knockout HEK cells devoid of any VRAC subunit^13^, so that the ectopically expressed LRRC8 proteins should not reach the cell surface (Fig. 3b). Indeed, it resulted in bright endomembrane fluorescence in transfected cells (not shown) but did not yield VRAC conductance (Fig. 3c). Only when we released functionally assembled VRACs formed by LRRC8A/E heteromers from the Golgi apparatus by UV-pulses, did we detect hypotonicity-induced VRAC conductance, but not in cells that were not illuminated with UV-pulses (Fig. 3c, d). Uncaged, plasma membrane delivered LRRC8A/E-composed VRACs showed the expected kinetics^13^ with maximal current observed ∼4 min after hypotonic shock (Supplementary Fig. 4). Furthermore, the recorded currents were completely blocked by DCPIB, the most selective inhibitor of VRAC^14,15^ (Fig. 3e). We concluded that we could restore ion channel function and thus osmotic regulation in knockout cells by releasing recombinant VRACs optogenetically.

**Figure 3.**
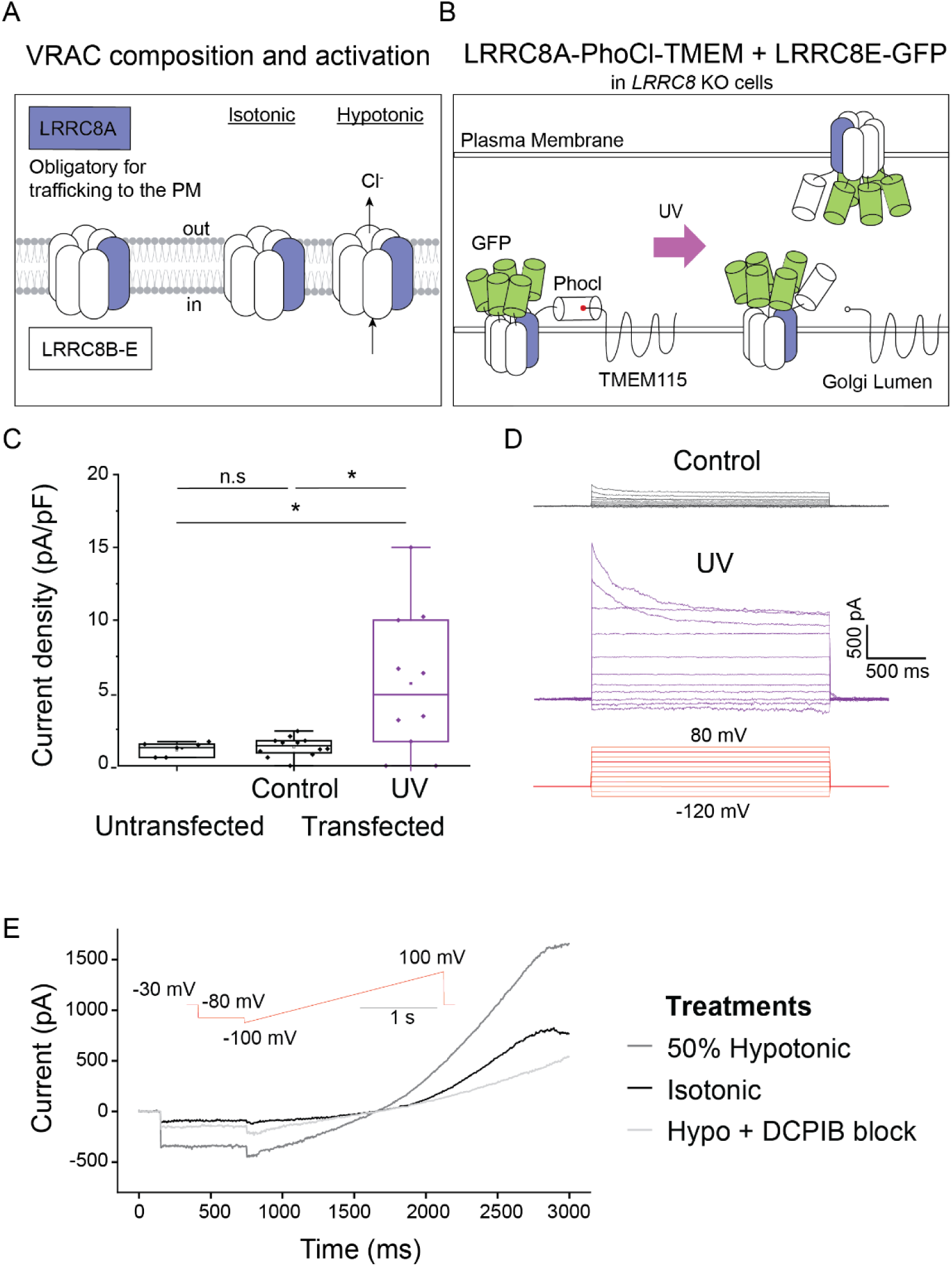
Optogenetic release of VRACs. **(a)** Composition (left) and activation of VRACs (right). **(b)** Schematic of VRAC’s release using PhoCl tool. VRACs composed of LRRC8A-PhoCl-TMEM and LRRC8E-GFP are expressed in *LRRC8* knockout cells. Following uncaging, VRACs traffic to the plasma membrane. **(c)** Quantification of current density at -80 mV after 50% hypotonic shock in untransfected HEK293T *LRRC8* knockout cells (mean, sd = 1 ± 0.4, n = 7, N = 1) and cells transfected with LRRC8A-PhoCl-TMEM and LRRC8E-GFP, without (control) (mean, sd = 1.3 ± 0.6, n = 12, N = 2) and with UV illumination (mean, sd = 5.6 ± 4.9, n = 10, N = 3). Significance was tested using the Mann-Whitney test (control vs UV, p = 0.03, untransfected vs UV, p = 0.04). **(d)** Current traces of activated VRAC measured using protocol shown in red in a control and a UV illuminated cell. **(e)** Time course of VRAC currents measured using voltage ramp protocol (shown in red) at isotonic, after 50% hypotonic shock and after DCPIB application in a representative UV illuminated cell.

Finally, we hypothesized that if a signaling pathway was disrupted by the knockout of an essential protein component of the system, the controlled release of the knocked-out component could restore the signaling event and allow for its quantitative investigation at the single cell level.

As a model, we used a T-lymphoblast system, in which the interleukin1 receptor (IL1R) is activated by its ligand in supported membrane bilayers after cell attachment. In these cells, IL1R-activation leads to the formation of a multi-protein signaling complex referred to as the Myddosome (Fig. 4a). We used a cell line, in which i) Myddosome protein MyD88 was genetically labeled with GFP and ii) Myddosome protein IRAK4, an essential kinase, knocked out via CRISPR/Cas9^16^. In these cells, activation of IL1R could not lead to downstream phosphorylation of inhibitor of nuclear factor kappa-B kinase (IKK) because of the lack of IRAK4^16^. When we now expressed IRAK4-mScarlet-PhoCl-TMEM at the Golgi apparatus in these cells and released IRAK4-mScarlet by a short pulse of 405 nm light, we found that IRAK4-mScarlet became recruited and co-assembled with MyD88 puncta on the plasma membrane as detected by TIRF microscopy (Fig. 4b, c). Furthermore, as previously shown^16^, IRAK4 regulates MyD88 oligomer size, and release of IRAK4 reduced MyD88 puncta size (Fig. 4d). Release of IRAK4 also induced the formation of functional Myddosome complexes leading to signal transduction. Indeed, when we fixed cells after optogenetic release of IRAK4-mScarlet and immunostained them for the phosphorylated form of inhibitor of nuclear factor kappa-B kinase subunit alpha and beta (pIKKα/β), the downstream output of Myddosome activation, we found that cells that contained MyD88 puncta colocalizing with the released IRAK4 were also positive for pIKKα/β staining, whereas Myddosomes without IRAK4 were not (Fig. 4e, f). We concluded that optogenetic release of the knocked-out IRAK4 could restore Myddosome-IKK signaling in single knockout T-Lymphoblasts.

**Figure 4:**
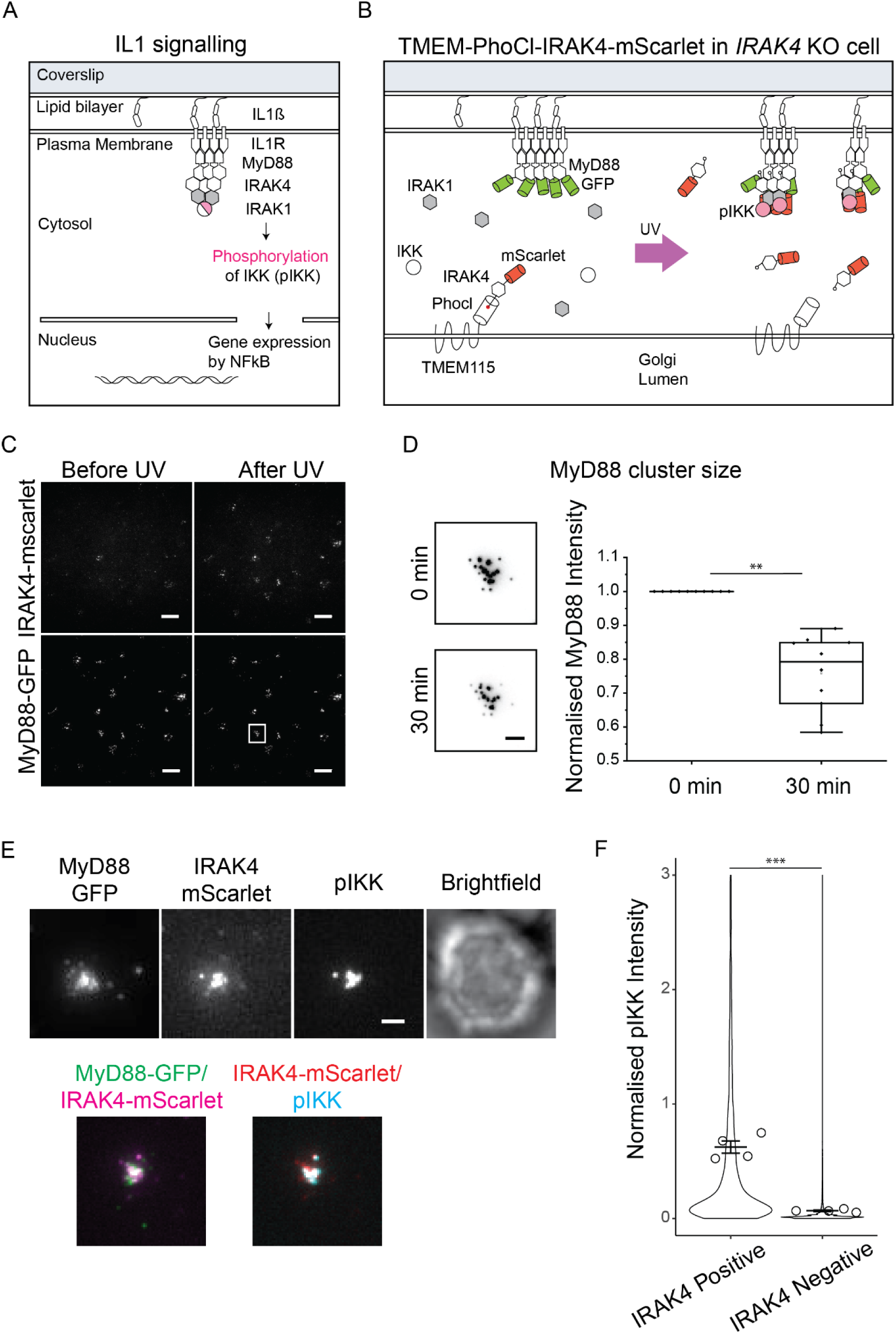
Reconstitution of IL1R signaling pathway in knockout cells with optogenetically released effector protein. **(a)** Schematic of IL1R signaling. **(b)** Schematic of TMEM-PhoCl-IRAK4-mScarlet construct in *IRAK4* knockout EL-4 cells. Without functional IRAK4 in the cells, the IL1R signaling is hampered and MyD88 forms big clusters. On release of IRAK4 with UV light, the IL1R signaling resumes and MyD88 cluster size is controlled. IL1R signaling results in phosphorylation of IKKα/β. **(c)** TIRF images of *IRAK4* knockout cells expressing MyD88-GFP and TMEM-PhoCl-IRAK4-mScarlet at 0 min and 30 min after UV illumination. Scale bars are 10 µm. **(d)** Left. Representative images of the same MyD88-GFP cluster at 0 min and 30 min after UV illumination. Scale bar is 2 µm. Right. Quantification of MyD88-GFP cluster size at 0 min (mean, sd = 1 ± 0) and 30 min after UV illumination (mean, sd = 0.75 ± 0.11) led IRAK4 release. Significance was tested using the Paired sample sign test (p = 0.0019). **(e)** TIRF images showcase IRAK4-mScarlet, MyD88-GFP and phosphorylated IKK-AF647 (pIKKα/β) cluster in *IRAK4* knockout cells expressing TMEM-PhoCl-IRAK4-mScarlet construct after optogenetic release of IRAK4 in the cytosol. Scale bar is 2 µm. **(f)** Quantification of pIKKα/β-AF647 intensity in *IRAK4* knockout cells without IRAK4 release (mean, sd = 0.6 ± 1, m = 2709) and after UV based IRAK4 release (mean, sd = 0.06 ± 0.01, m = 112242). The violin plot shows the distribution of individual MyD88 puncta measurements. Dots superimposed on the violin plot represent means of replicates (N = 4). Bars represent mean ± SEM. Significance was tested using the two-tailed unpaired t-test (p = 10^-5^), employing the means of replicates as data points. m = number of MyD88 puncta analyzed. SEM = Standard Error of the Mean.

## Discussion

We demonstrate here an optogenetic system for the release of single functional molecules in live cells. Our approach overcomes an important bottleneck in the observation of single molecules in that it allows for the controlled release of small amounts of functional protein, compatible with the density requirements of single molecule imaging. All cytosolic and membrane proteins should be compatible with our system as we have N-terminal and C-terminal anchors for the Golgi apparatus and even functional multispanning and multidomain transmembrane complexes such as BK and VRAC ion channels could be optogenetically released. Furthermore, our systems achieve quantitative labeling efficiency, thus giving access to single molecule imaging of any molecule without cellular background that may interfere in the isolated observation of specific function. We thus generated a straightforward approach for single molecule imaging of any membrane or cytosolic protein in live cells.

Optogenetic release of single molecules has the potential to address previously intractable questions regarding the numbers and dynamics of proteins required to execute cellular functions. In this way, the precise molecular function of many proteins could be isolated using a combination of knockout cell lines with quantitative delivery of the depleted molecule via optogenetic release. We here demonstrate this capability showing that the optogenetic release of IRAK4 in knockout cells reconstitutes Myddosome formation, IKK activation in IL-1 signaling. In the future, our technique may allow determination of the number of activated molecules that are required to reach a tipping point leading to a functional decision at the level of the entire cell in similar systems.

We demonstrate the reconstitution of specific ion channel conductance in cells otherwise devoid of them (Fig. 2, 3). The high sensitivity of electrophysiology as well as our results from single molecule imaging of released molecules (Fig. 2) demonstrate the extremely low leakiness of our method.

Releasing a controlled wave of functional, tagged proteins in live cells for acute investigation is highly advantageous. In contrast, when a cell expresses a labeled protein of interest in steady-state, the population of target proteins could be heterogenous in terms of its conformation, post-translational modifications and interaction with binding partners. In such a situation, any observation of a specific type of process will thus be biased by the preexisting distribution of states of the molecule. As a result, the measurement will experience a high level of noise. Our method, by releasing a small wave of a target molecule, allows for a synchronized observation of the sequence of events that the target molecule is involved in as it executes its function.

In the future, our method could be further improved in terms of photoconversion efficiency and dissociation rate. PhoCl is the first reported photocleavable fluorescent protein so far, but many molecules could be engineered to generate future variants. The mechanism by which PhoCl was generated from mMaple^17^, circular permutation and mutagenesis, should in principle be applicable to all fluorescent proteins that experience peptide backbone breakage after UV illumination such as the mEOS proteins^18^, Kaede^19^, mkikGR^20^, Dendra2^21^ or IrisFP^22^. While we here already used an improved variant of PhoCl^23^ with faster release kinetics of the cleaved peptide, we expect more and further improved photocleavable proteins will be generated in the future, extending the range and capabilities of our concept. Finally, to extend the scope of our method from single molecule imaging and functional reconstitution in live cells, it may be possible to release proteins via PhoCl in animals via 2-photon mediated photocleavage as well, allowing to express proteins in specific cells at a specific time inside developing embryos or living animals. In this context, precisely timed release of functional protein during organism development or behavior in a knockout background should generate unprecedented resolution of molecular processes in complex, dynamic environments.

## Materials and Methods

### Generation of constructs

For TMEM-PhoCl-FGF-GFP, the cDNA encoding for human TMEM115 (Origene cat# RG203956) was cloned into pEGFP-C1 using NheI and BsrGI restriction sites. PhoCl, amplified from pcDNA-NLS-PhoCl-mCherry, was cloned into pEGFP-TMEM115 plasmid using XhoI and HindIII restriction sites. Finally, FGF2-GFP was cloned into pEGFP-TMEM115-PhoCl plasmid using EcoRI and SalI restriction sites. pcDNA-NLS-PhoCl-mCherry is Addgene plasmid #87691; http://n2t.net/addgene:87691; RRID: Addgene_87691). In all further work, a novel, improved variant of PhoCl called PhoCl-2c^24^ was used. From here on, PhoCl will stand for PhoCl-2c.

For mScarlet-CD4-PhoCl-RER, fusion construct encompassing a signal peptide (MWPLVAALLLGSACCGSA), genes for mScarlet, the transmembrane region of CD4 (STPVQPMALIVLGGVAGLLLFIGLGIFFCVRCRHRRR), PhoCl and RER1 was designed and generated by gene synthesis (Thermo Fisher Scientific). The fusion construct was then cloned into pTREtight2 vector (Addgene) using NheI and XbaI (New England Biolabs). pTREtight2 was a gift from Markus Ralser (Addgene plasmid #19407; http://n2t.net/addgene:19407; RRID: Addgene_19407).

To generate the FLAG-BK-YFP-PhoCl-TMEM construct, the Gibson cloning method^25,26^ was used. Briefly, the fragment for the FLAG-BK-YFP, PhoCl-2c and TMEM115 were cloned into the pTREtight2-vector plasmid using NEBuilder HiFi DNA Assembly (New England Biolabs). The FLAG-BK-YFP fragment was amplified from FLAG-BK667YFP plasmid^27^, a gift from Teresa Giraldez.

LRRC8A-PhoCl-TMEM construct was generated from LRRC8A-GFP-PhoCl-TMEM plasmid. The LRRC8A-GFP-PhoCl-TMEM plasmid was digested with AgeI (NEB) and EcoRI (NEB) to produce three fragments, LRRC8A fragment, GFP fragment and the remaining vector with PhoCl-TMEM115 genes (PhoCl-TMEM-vector). The LRRC8A fragment was fused to the PhoCl-TMEM-vector fragment by the Gibson cloning method. The LRRC8A-GFP-PhoCl-TMEM plasmid used as a template was also constructed via the Gibson cloning method, where fragments encoding LRRC8A, GFP, PhoCl, and TMEM115 were fused in pTREtight2 vector.

For the TMEM-PhoCl-IRAK4-mScarlet construct, TMEM-PhoCl with a fragment with a 3xGGS linker was generated using PCR from TMEM-PhoCl-rtTA3-mScarlet, a plasmid synthesized and cloned by Twist Bioscience. IRAK4, ordered as a gBlock (IDT), was fused to mScarlet via a 3xGGS linker by PCR. A vector backbone was generated using pHR-dSV lentiviral plasmid digested with Mlu1 and Not1 restriction enzymes (New England Biolabs). Construction of the pHR-dSV-TMEM-PhoCl-IRAK4-mScarlet fusion was achieved by Gibson Assembly of the three components.

### Cell culture and transfection

CV1 cells, used for imaging secreted CD4 transmembrane proteins and BK-channels, were maintained with high-glucose, indicator-free Dulbecco’s Modified Eagle Medium (DMEM) (Thermo Fisher Scientific) supplemented with 10 % FBS (Thermo Fisher Scientific) and 1 % glutaMax (Thermo Fisher Scientific). CHO-K1 cells stably expressing TMEM-PhoCl-FGF2-GFP cells were cultured in Minimum Essential Medium (MEM)-α medium (Sigma-Aldrich) supplemented similarly with 10 % FBS and 1 % glutaMax. Gene-edited EL-4 cells, used for Mydosome signaling assays, were cultured in RPMI (Thermo Fisher Scientific) with 10 % FBS supplemented with 1% glutamine. For BK patch-clamp experiments, HEK293T cells were cultured in MEM (PAN-Biotech) supplemented with 6 % FBS, 1 % penicillin/streptomycin and 1 % glutamine. For VRAC patch-clamp experiments, LRRC8 knock-out HEK293T cells (disrupted of all five *LRRC8* genes)^13^ were cultured in DMEM (PAN-Biotech) supplemented with 10 % FBS and 1 % glutamate. All cells were maintained at 37 °C with 5 % CO_2_.

Fusion constructs with pTREtight2 vectors were co-transfected with reverse tetracycline-controlled transactivator 3 (pLenti CMV rtTA3 Hygro (w785-1)) plasmid (Addgene) in the presence of doxycycline (Clontech). pLenti CMV rtTA3 Hygro (w785-1) was a gift from Eric Campeau (Addgene plasmid #26730; http://n2t.net/addgene:26730; RRID: Addgene_26730). pLenti CMV rtTA3 Hygro is called rtTA3 in the next sections.

### Generation of stable cell line

CHO-K1 cells stably expressing TMEM-PhoCl-FGF2-GFP in a doxycycline-dependent manner were generated using a retroviral transduction system based on Moloney Murine Leukemia Virus, as previously described^28^. In brief, for recombinant virus particle production, HEK 2-293 cells were used, which contained the retroviral packaging proteins pVPack-GP and pVPack-eco (HEK EcoPack 2-293 cells). These cells were transfected with TMEM-PhoCl-FGF2-GFP within the pRev-TRE2 vector, containing a TET-Responsive Element (TRE2). Cell transfection and virus production was performed using the MBS Mammalian Transfection Kit (Agilent Technologies). The infectious supernatant was collected 72 hours after transfection and filtered onto CHO-K1 target cells for transduction. Transduced cells stably expressed the Murine Cationic Aminoacid Transporter (MCAT-1) and rtTA2-M2, a Tet-ON transactivator. Following 72 hours of incubation with the virus, cells were subjected to three subsequent FACS sorting versus GFP fluorescence. The second sorting step was conducted in the absence of doxycycline to select cells not expressing the protein in the absence of induction. The first and the last sorting steps were conducted following 24 hours of incubation with doxycycline to select expressing cells.

For generation of EL-4 cells stably expressing TMEM-PhoCl-IRAK4-mScarlet, co-transfection of pHR transfer plasmids with second-generation packaging plasmids pMD2.G and psPAX2 (a gift from Didier Trono, Addgene plasmid #12259 and #12260) was done to produce lentivirus in HEK293T cells. Virus particles were harvested from the supernatant after 72 hours, filtered and applied to *IRAK4* knockout EL-4 cells. Cells were then FACS sorted to select populations positive for the pHR-dSV-TMEM-PhoCl-IRAK4-mScarlet. EL4.NOB1 cells were used as a negative control for FACS gating. Cells were sorted using a BD FACS Aria II at the Deutsches Rheuma-Forschungszentrum Berlin Flow Cytometry Core Facility.

### Live cell microscopy-

PhoCl activation and standard fluorescence microscopy were performed on an inverted Olympus IX71 microscope equipped with a Yokogawa CSU-X1 spinning disk; A 60x/ 1.42 NA oil Olympus objective was used together with 491 nm (100 mW; Cobolt), 561 nm (100 mW; Cobolt) laser. A quad-edge dichroic beam splitter (446/ 523/ 600/ 677 nm; Semrock) was used to separate fluorescence emission from excitation light, and final images were taken with an Orca Flash 4.0 sCMOS camera (Hamamatsu). Images were acquired with MetaMorph (Molecular Devices).

All TIRF single molecule tracking experiments were performed on a custom-built microscope^29^. An Olympus TIRF objective (60x/ 1.49 NA) was used with a 473 nm laser (100 mW; Laserglow Technologies) and a 643 nm laser (150 mW; Toptica Photonics). A quad-edge dichroic beamsplitter (405/ 488/ 561/ 635 nm; Semrock) separated fluorescence emission from excitation light. The emission light was further filtered by a quad-band bandpass filter (446/ 523/ 600/ 677 nm; Semrock) and focused by a 500 mm tube lens onto the chip of a back-illuminated electron-multiplying charge-coupled device camera (Evolve; Photometrics) that was water-cooled to -85 °C. Images were acquired with MicroManager^30^.

### CD4 and BK pulse chase assay

CV1 cells were transfected with mScarlet-CD4-PhoCl-RER and rtTA3 plasmids using the Neon Transfection System (Thermo Fisher Scientific) 24 hours before the experiment. They were seeded onto glass-bottom 35-mm gridded dishes (Ibidi) in the presence of doxycycline (0.02 µg/ml). Plasma membrane-localized mScarlet-CD4 was detected using RFP antibody. Before uncaging, cells were incubated with RFP antibody (Chromotek) labelled with NHS AF647 for 15 min and subsequently imaged in TIRF microscopy. For uncaging, cells were illuminated with 5 s pulses of a 405 nm laser at 33 mW/mm^2^ every 15 s for five times. Three hours after the uncaging, the cells were again incubated with anti-RFP antibodies coupled to AF647 for 15 min and subsequently imaged in TIRF microscopy to detect surface-delivered mScarlet-CD4. The TIRF images of secreted mScarlet-CD4 molecules on the plasma membrane were analyzed using ImageJ. Particles were detected and counted using the Trackmate^31^ plugin in ImageJ^32^. The number of particles was normalized to the cell surface area and time. Pulse-chase assay for BK-channels was performed similarly. FLAG-BK-YFP-PhoCl-TMEM was co-transfected with rtTA3 in the presence of doxycycline (1µg/ml). FLAG antibody (Sigma Aldrich) labelled with NHS AF647 was used to detect the secreted BK-channels. Secreted BK-channels were detected seven hours after the uncaging step.

### TMEM-PhoCl-FGF2-GFP activation and uncaged FGF2 quantification

CHO-K1 cells stably expressing TMEM-PhoCl-FGF2-GFP were seeded on round #1, 18 mm diameter glass coverslips. Cells were illuminated with 5 s pulses of a 405 nm laser at 33 mW/mm^2^ every 15 s for five times for PhoCl photoconversion and cleavage. Images were acquired before and 10 min after the 405 nm illumination. Mean intensity within the cell boundary was measured using ImageJ to quantify cytosolic FGF2-GFP release after uncaging.

For quantifying membrane bound FGF2 after uncaging, a TIRF microscopy assay was performed. Cells were illuminated with 10 s pulses of a 405 nm laser at 2 mW/mm^2^ for uncaging. Movies were acquired before and after the 405 nm illumination at TIRF mode in 491 nm channel to image FGF2-GFP. The number of plasma membrane-bound FGF2-GFP molecules was measured by tracking the particles using the Trackmate plugin in ImageJ. The number of particles was normalized by the cell surface area and time.

### IRAK4 reconstitution assay

Imaging of EL-4 cells was performed on an inverted microscope (Nikon TiE) equipped with a Nikon fiber launch TIRF illuminator. Illumination was controlled with a laser combiner using the 488 nm, 561 nm, and 640 nm laser lines. Fluorescence emission was collected through filters for GFP (525 ± 25 nm), RFP (595 ± 25 nm), and AF647 (700 ± 75 nm). All images were collected using a Nikon Plan Apo 100x 1.4 NA oil immersion objective that projected onto a Photometrics 95B Prime sCMOS camera with 2 × 2 binning (calculated pixel size of 150 nm) and a 1.5x magnifying lens. Image acquisition was performed using NIS-Elements software. Live cell experiments were performed at 37°C. The microscope stage temperature was maintained using an OKO Labs heated microscope enclosure. To release PhoCl-IRAK4, EL-4 cells were illuminated with a 405 nm laser line for 10 s at 21 mW/mm^2^ laser power. After uncaging, TIRF images of MyD88-GFP and IRAK4-mScarlet-PhoCl-TMEM were acquired for 30 min with an interval of 30 s.

### IL-1β-functionalized supported lipid bilayers

Supported lipid bilayers (SLBs) were prepared using a previously published method^16^. Briefly, Phospholipid mixtures consisting of 97.5 % mol 1-palmitoyl-2-oleoyl-*sn*-glycero-3-phosphocholine (POPC), 2 % mol 1,2-dioleoyl-sn-glycero-3-[(N-(5-amino-1-carboxypentyl)iminodiacetic acid)succinyl] (ammonium salt) (DGS-NTA) and 0.5 % mol 1,2-dioleoyl-sn-glycero-3-phosphoethanolamine-N-[methoxy(polyethylene glycol)-5000] (PE-PEG5000) were mixed in glass round-bottom flasks and dried down with a rotary evaporator. All lipids used were purchased from Avanti Polar Lipids. Dried lipids were placed under vacuum for 2 hours to remove trace chloroform and resuspended in PBS. Small unilamellar vesicles (SUVs) were produced by several freeze-thaw cycles combined with bath sonication. Once the suspension had cleared, the SUVs were spun in a benchtop ultracentrifuge at 35,000xg for 45 min. SUVs were stored at 4°C for up to a week.

To prepare SLBs on 96-well glass bottom plates, the plates were cleaned for 30 min with a 5% Hellmanex solution containing 10% isopropanol heated to 50°C, then incubated with 5% Hellmanex solution for 1 hour at 50°C, followed by extensive washing with distilled water. 96-well plates were dried with nitrogen gas and sealed until needed. To prepare SLBs, individual wells were cut out and base etched for 15 min with 5 M KOH and then washed with PBS. To form SLBs, SUVs suspension was deposited in each well or coverslip and allowed to form for 1 hour at 45°C. After 1 hour, wells were washed extensively with PBS. SLBs were incubated for 15 min with HEPES buffered saline (HBS: 20 mM HEPES, 135 mM NaCl, 4 mM KCl, 10 mM glucose, 1 mM CaCl_2_, 0.5 mM MgCl_2_) with 10 mM NiCl_2_ to charge the DGS-NTA lipid with nickel. The SLBs were then washed in HBS containing 0.1% BSA to block the surface and minimize non-specific protein adsorption. After blocking, the SLBs were functionalized by incubation with His10-IL-1β for 1 hour. The labeling solution was washed out and each well was filled with HBS with 0.1% BSA. For SLBs set up on 96-well plates the total well volume was 69 μl, and 57 μl was removed leaving 120 μl of HBS 0.1% BSA in each well. Each SLBs was functionalized with 120 μl His10-Halo-IL-1β of twofold desired concentration for 1 hour and excessive ligands were washed away with HBS.

### Immunofluorescence staining

To analyze the downstream activation of phospho-IKKα/β on MyD88-GFP puncta, EL-4 cells were stimulated with IL-1β-functionalized SLBs for 30 min, followed by 405 nm illumination (10 s at 21 mW/mm^2^) to release PhoCl-IRAK4. After 45 min, cells were fixed with 3.5 % PFA containing 0.5 % Triton X-100 for 20 min at room temperature. Cells were washed with PBS and blocked with PBS 10 % BSA at 4°C overnight. The next day, cells were incubated with an antibody mixture containing anti-phospho-IKKα/β (Cell Signaling) and Nano-secondary AF647 (Chromotek) prepared in PBS with 10 % BSA and 0.1 % Triton X-100 for 1 hour at room temperature. Cells were then labeled with FluoTag-X4 anti-GFP conjugated to Atto488 (NanoTag Biotechnologies) and FluoTag-X2 anti-mScarlet conjugated to Atto565 (NanoTag Biotechnologies) at room temperature for 1 hour. After antibody labelling, cells were washed with PBS and imaged with TIRF microscopy.

### Quantification and analysis of MyD88-GFP puncta

To quantify the intensity of MyD88-GFP puncta, images were processed in ImageJ to remove background fluorescence using custom-written macros described previously^16^. Briefly, the processing step included image subtraction with a dark frame image to remove camera noise, followed by image subtraction with a median-filtered image to remove the background associated with cytosolic fluorescence. The dark frame image was acquired with no light exposure to the camera but identical exposure time to experimental acquisition while the median-filtered image was generated in Fiji with a radius of 25 pixels. Post processing, individual MyD88-GFP puncta were manually selected, and their mean intensities were measured. MyD88-GFP puncta without IRAK4 association were used as reference for correction of photobleaching.

### Quantification and analysis of pIKKα/β staining

TIRF microscopy images of pIKKα/β immunofluorescence staining was quantified using a previously described analysis pipeline^33^. First, the dark frame and median-filtered images were subtracted from the MyD88-GFP and pIKKα/β staining TIRF micrographs. Next, MyD88-GFP puncta were segmented, and their fluorescence intensity was measured using a custom Cell Profiler pipeline. The integrated intensity and mean intensity of the MyD88, IRAK4 and pIKKα/β were extracted for each segmented puncta. Data normalization and visualization were performed using R. Intensity normalization was performed using the following equation:

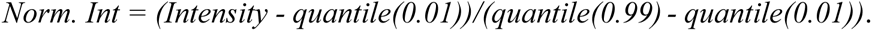

MyD88 puncta was defined as IRAK4 positive if IRAK4 normalized integrated intensity equal to or exceeded 0.5. pIKKα/β normalized integrated intensity was plotted.

### BK-channel patch clamp assay

HEK293T cells transfected with FLAG-BK-YFP-PhoCl-TMEM and rtTA3 were seeded on glass-bottom 35-mm gridded dishes in the presence of doxycycline (1µg/ml). Whole-cell patch-clamp recordings were performed at room temperature one day after transfection and doxycycline induction. Cells were illuminated with 15 s pulses of 405 nm at 1.42 mW/mm^2^. Experiments were carried out 7 hours after UV exposure (uncaged cells) and control measurements were simultaneously performed on separate dishes (non-uncaged cells).

Currents were acquired using an Axopatch 200B amplifier and the Axograph acquisition program (Axograph Scientific) via an Instrutech ITC-18 D-A interface (HEKA Elektronik). Currents were filtered at 5 kHz and digitized at 20 kHz. Bath solution was composed as follows, 145 mM NaCl, 5 mM KCl, 1 mM MgCl_2_, 10 mM HEPES, 3 mM EGTA, pH 7.4. Given the high conductance of BK-channels the following internal (pipette) solution was used to reduce the current and minimize the series resistance error, 90 mM NMDG-Cl, 50 mM KCl, 4 mM NaCl, 10 mM HEPES, 1 mM Mg-ATP, 10 mM EGTA, 2 mM CaCl_2_ pH 7.3^34,35^. Pipette resistance was between 2 and 6 MΩ, and the cell capacitance was between 3 and 25 pF, as measured by the compensating circuit of the amplifier. We accepted a maximal voltage-clamp error (calculated as the product of the maximum current and the uncompensated series resistance) of 10 mV. The standard IV-protocol to elicit BK currents consisted of 100 ms voltage steps ranging from -80 mV to +140 mV in 20 mV increments, starting from a holding potential of 0 mV. No leak-current subtraction was performed. Figures were prepared using Igor Pro (https://www.wavemetrics.com/products/igorpro).

### VRAC patch clamp assay

*LRRC8* knock-out HEK293T cells (disrupted of all five *LRRC8* genes)^13^ were plated onto glass-bottom 35-mm gridded dishes and transfected using the Ca_3_(PO_4_)_2_ technique. The transfected LRRC8E was fused C-terminally with GFP. LRRC8E-GFP and LRRC8A-PhoCl-TMEM were co-transfected with rtTA3 in presence of doxycycline (1µg/ml). Cells were illuminated with 10 s pulses of a 405 nm light (58 mW/mm^2^) (CoolLED pE-300ultra multi-band spectrum) for PhoCl photoconversion and cleavage. Experiments were carried out 4 hours after UV exposure (uncaged cells) and control measurements were simultaneously performed on separate dishes (non-transfected and non-uncaged cells).

Whole-cell voltage-clamp experiments were performed in isotonic extracellular solution containing 150 mM NaCl, 6 mM KCl, 1 mM MgCl_2_, 1.5 mM CaCl_2_, 10 mM glucose, and 10 mM HEPES, pH 7.4 with NaOH (320 mOsm). VRAC currents were elicited by perfusing the cells with hypotonic solution containing 75 mM NaCl, 6 mM KCl, 1 mM MgCl_2_, 1.5 mM CaCl_2_, 10 mM glucose, 10 mM HEPES, pH 7.4 with NaOH (160 mOsm). The pipette solution contained 40 mM CsCl, 100 mM Cs-methanesulfonate, 1 mM MgCl_2_, 1.9 mM CaCl_2_, 5 mM EGTA, 4 mM Na_2_ATP, and 10 mM HEPES, pH 7.2 with CsOH (290 mOsm). Osmolarities of all solutions were assessed with an Osmometer OM 807 freezing point osmometer (Vogel). All experiments were performed at a constant temperature of 20–22°C. Currents were recorded with an EPC-10 USB patch-clamp amplifier and PatchMaster software (HEKA Elektronik). Patch pipettes had a resistance of 3–5 MΩ. Currents were sampled at 5 kHz and low-pass filtered at 10 kHz. The holding potential was -30 mV. The standard protocol for measuring the time course of VRAC current activation consisted of a 0.6 s step to -80 mV followed by a 2.6 s ramp from -100 to 100 mV and was applied every 12 s. The read-out for VRAC current was the steady-state whole-cell current at -80 mV normalized to the cell capacitance (current density) subtracted by the baseline current density at -80 mV before application of hypotonic solution. The voltage protocol, applied before the standard protocol and after complete activation of VRAC, consisted of 2 s steps starting from -120 mV to 80 mV with 20-mV increment preceded and followed by a 0.5 s step to -80 mV every 5 s. The voltage-step protocol confirmed VRAC-typical properties of outward rectification and depolarization-dependent inactivation for LRRC8A/E-containing VRAC^13^. In the uncaged cells VRAC currents were blocked by 100 μM DCPIB (4-(2-butyl-6,7-dichloro-2-cyclopentyl-indan-1-on-5-yl) oxybutyric acid, Tocris).

## Author contributions

H.E. and P.K. conceived of and designed the project. P.K., X.L., R.C., F.B., N.S., W.N. and R.S. contributed reagents, D.B. helped in initial stages of the project. P.K. and F.B performed and analyzed all single molecule imaging experiments. S.B. and A.P. designed, performed and analyzed the electrophysiology experiments for BK-channels. Y.K. and T.S. designed, performed and analyzed the electrophysiology experiments for VRAC. F.C., N.S. and M.J.T. designed, performed and analyzed the experiments with IRAK4. H.E. and P.K. wrote the manuscript with input from all authors.

## Acknowledgements

We thank Teresa Giraldez for the kind gift of the plasmid encoding the BK-channel, Slo1. This work was funded by the Deutsche Forschungsgemeinschaft (DFG, German Research Foundation) as part of TRR 186 (Project Number 278001972), and under Germanýs Excellence Strategy -EXC-2049-390688087 “NeuroCure”. A.P. was recipient of a Heisenberg Professorship from the DFG (Projektnummer 446182550). Work in the lab of R.E.C. was supported by grants from the Canadian Institutes of Health Research (FS-154310) and the Natural Sciences and Engineering Research Council of Canada (RGPIN-2018-04364) The authors would like to thank Anna Klemmer, Carolin Knappe, Claire Schlack, Daria Shyshko, Andrea Senge and Aurora Elhazaz Fernandez for help with molecular biology and all members of the Ewers laboratory for helpful discussions.

## Supplementary Figures

**Supplementary Figure 1.**
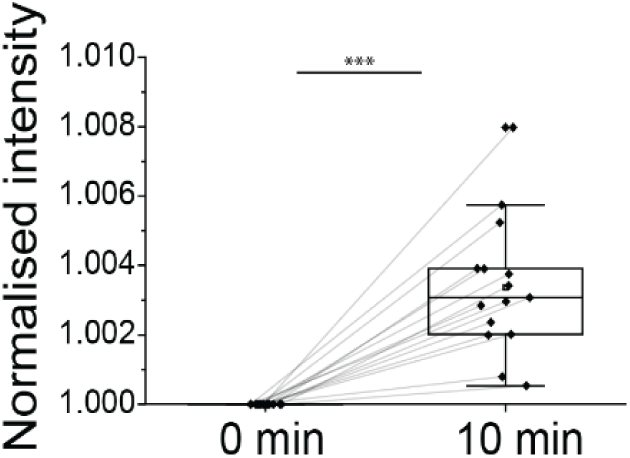
Release of FGF2-GFP in the cytosol. Quantification of cytosolic FGF2-GFP before (mean, sd = 1 ± 0) and 10 min after UV Illumination (mean, sd = 1.05 ± 0.02) of CHO cells transiently expressing TMEM-PhoCl-FGF2-GFP, N = 1, n = 14. Significance was tested using the Paired sample sign test (p = 10^-4^).

**Supplementary Figure 2.**
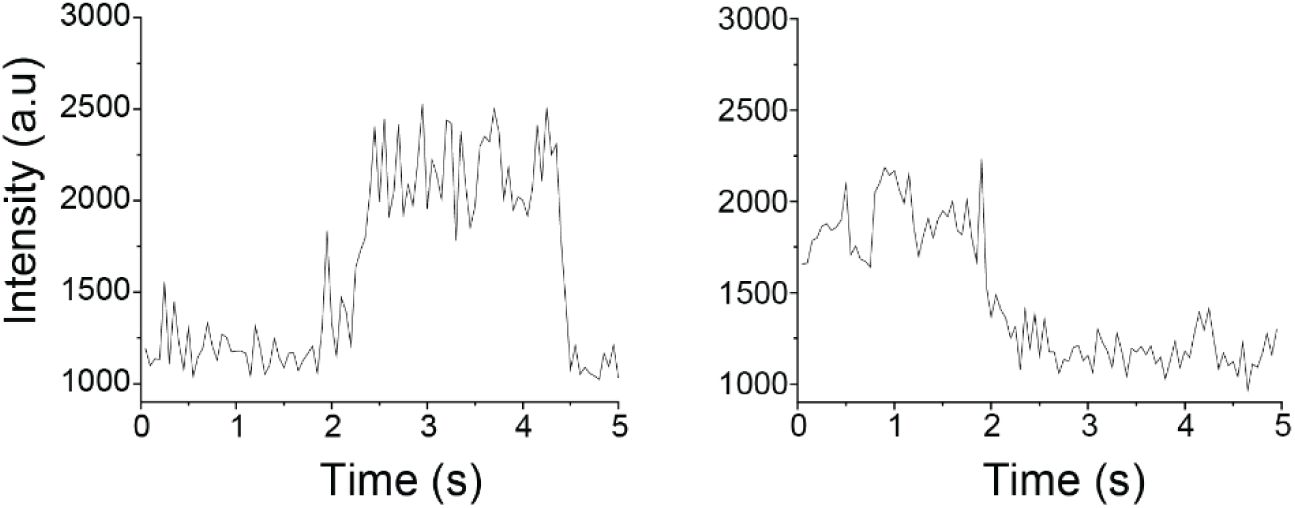
Single FGF2-GFP at the plasma membrane. Time traces of fluorescence emission from FGF2-GFP molecules on the plasma membrane.

**Supplementary Figure 3.**
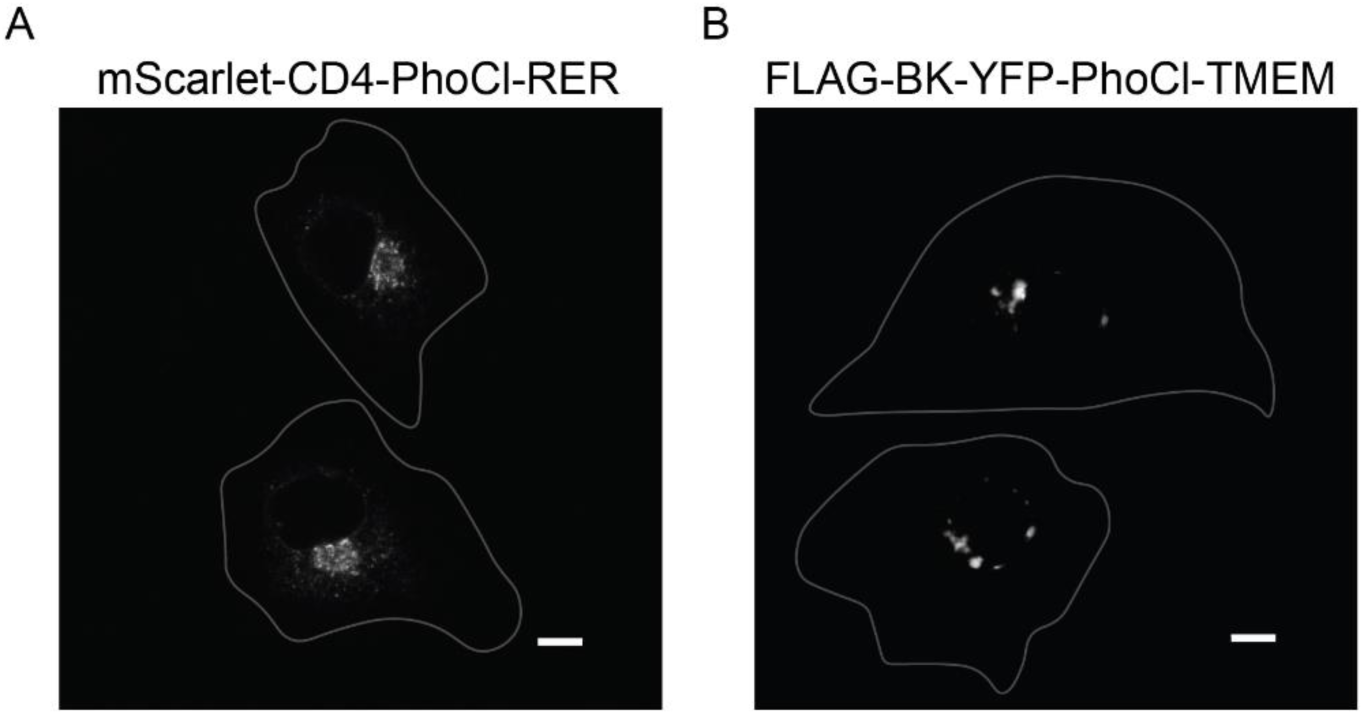
Golgi localization of PhoCl caged proteins. **(a)** CV1 cells expressing mScarlet-CD4-PhoCl-RER (b) CV1 cells expressing FLAG-BK-YFP-PhoCl-TMEM. Scale bars are 10 µm.

**Supplementary Figure 4.**
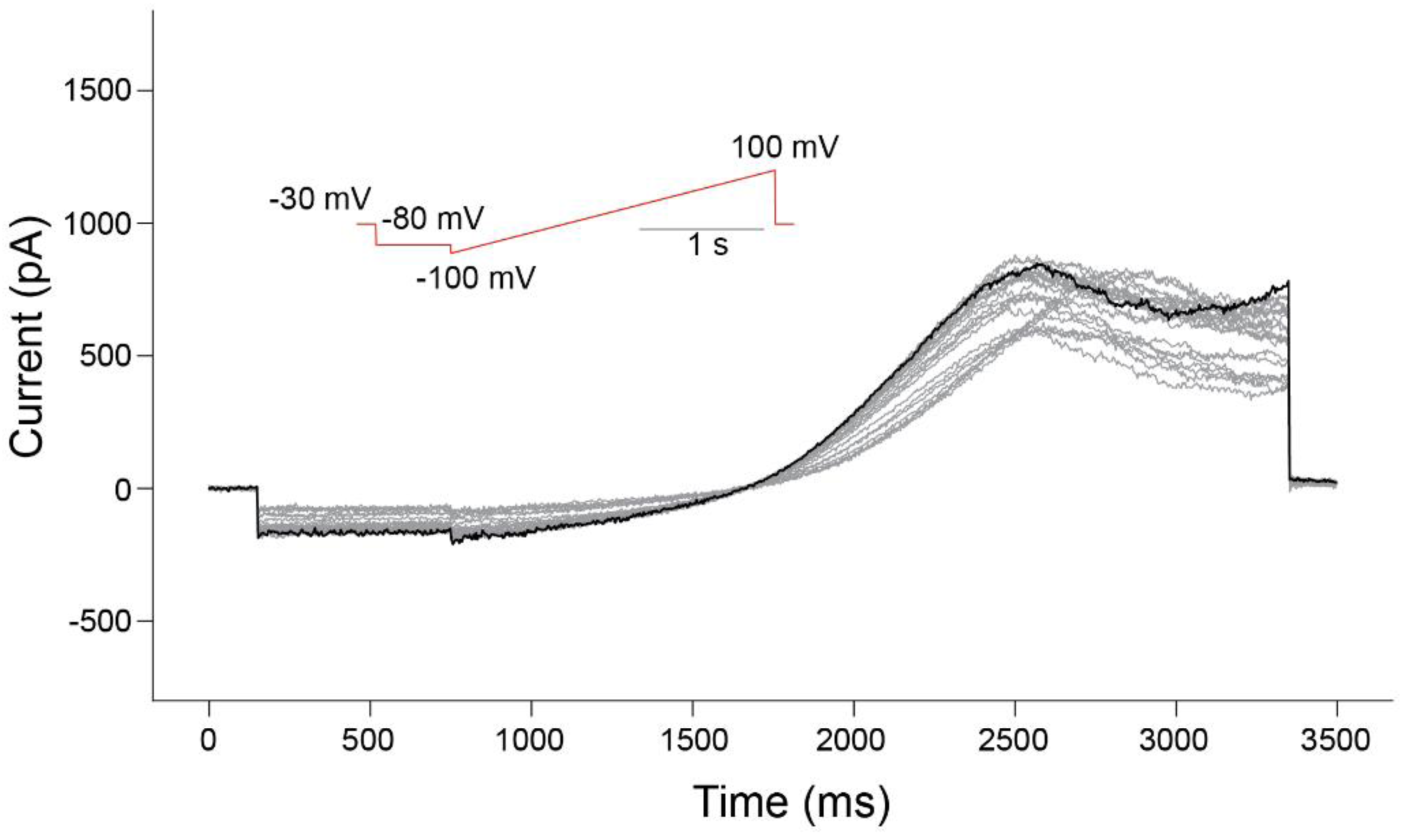
Current kinetics of optogenetically released VRAC. Example ramp current traces from optogenetically released VRACs for 4 min upon switching bath solution to hypotonicity with maximal developed current trace shown in black. Ramp protocol shown red in inset. Note fast inactivation at inside-positive voltages typical for LRRC8A/E-composed VRACs.

